# Single-cell profiling of histone modifications in the mouse brain

**DOI:** 10.1101/2020.09.02.279703

**Authors:** Marek Bartosovic, Mukund Kabbe, Gonçalo Castelo-Branco

## Abstract

The development of the mouse central nervous system (CNS) involves coordinated execution of transcriptional and epigenetic programs. These programs have been extensively studied through single-cell technologies in a pursuit to characterize the underlying cell heterogeneity. However, histone modifications pose additional layers of both positive and negative regulation that defines cellular identity. Here we show that the Cut&Tag technology can be coupled with a droplet-based single cell library preparation platform to produce high quality chromatin modifications data at a single cell resolution in tens of thousands of cells. We apply single-cell Cut&Tag (scC&T) to probe histone modifications characteristic of active promoters (H3K4me3), active promoters and enhancers (H3K27ac), active gene bodies (H3K36me3) and inactive regions (H3K27me3) and generate scC&T profiles for almost 50,000 cells. scC&T profiles of each of these histone modifications were sufficient to determine cell identity and deconvolute at single cell level regulatory principles such as promoter bivalency, spreading of H3K4me3 and promoter-enhancer connectivity. Moreover, we used scC&T to investigate the single-cell chromatin occupancy of transcription factor Olig2 and the cohesin complex component Rad21. Our results indicate that analysis of histone modifications and transcription factor occupancy at a single cell resolution can provide unique insights of epigenomic landscapes in the CNS. We also provide an online resource that can be used to interactively explore the data at https://castelobranco.shinyapps.io/BrainCutAndTag2020/.

## Main text

For decades, chromatin immunoprecipitation (ChIP) coupled with deep sequencing (ChiP-seq) has been the gold standard in the field of epigenetics. However, ChIP-seq experiments inherently suffer from low signal-to-noise ratio, inconsistencies due to immunoprecipitation and high requirements for sample quantity. Recently, novel methods based on *in situ* chromatin cleavage or tagmentation have been introduced. In particular, the development of Cut&Run^1^ and Cut&Tag technologies^2^, with low input requirements, have put forward the possibility of investigating chromatin modifications at the single cell level in large scale^2^.

The onset of single-cell sequencing technologies marked a new era in developmental biology. It has allowed exploratory analysis of tissue complexity and cell heterogeneity^3^. The wave of exploratory studies was followed by more in-depth analysis of gene regulatory networks, imputation of developmental trajectories and prediction of the future cell states (e.g. RNA velocity)^4^. Moreover, technologies that focus on probing the epigenetic landscape such as single-cell ATAC-seq^5^ or single-cell DNA methylation^6^ sequencing have also emerged, shedding light on the epigenetic heterogeneity of tissues. While chromatin accessibility and DNA methylation can provide genome-wide snapshots of active and repressive states, the study of diverse chromatin modifications might provide further insights on epigenomic and cellular states.

Modifications of histone tails represent a unique system of regulation of gene expression. Histone acetyltransferases and methyltransferases deposit post-translational modifications at various genomic elements (e.g. promoters, enhancers) to regulate gene expression both positively (e.g. H3K27ac, H3K4me3, H3K4me1) and negatively (e.g. H3K27me3, H3K9me3). Bulk studies of these modifications have succeeded in defining the regulatory elements but failed to uncover potential cell heterogeneity within tissue samples. Recent studies have characterized the state of post-translational modifications of histones at a single cell level in cultured cells and embryos^2,7–11^. However, for highly complex adult organs such as the brain, this single-cell characterization has not been achieved.

Here, we developed and applied single-cell Cut&Tag protocol by adapting the droplet-based 10x Genomics single cell ATAC-seq platform, to investigate histone modification profiles at the single cell level in the mouse brain. We focused on the oligodendrocyte lineage (OLG), which we have recently shown to be heterogeneous and to be able to transition to alternative cell states during development and disease^12–15^. We were able to resolve single cells into discrete populations based exclusively on the histone modifications data, find unique cell type specific markers and quantitative differences in the levels of histone modifications. We used the obtained data to get unique insights into promoter mark spreading, bivalency and identification of enhancer-promoter interactions. Finally, we were able to obtain single cell binding profiles for non-histone proteins, namely chromatin architecture factor and subunit of cohesin complex Rad21 and the OLG-specific transcription factor Olig2. We have generated a web resource available at https://castelobranco.shinyapps.io/BrainCutAndTag2020/, where these scC&T datasets can be explored. Altogether this study provides a novel perspective on epigenetic heterogeneity within the brain and opens up possibilities to study epigenetic regulation in complex tissues in great detail.

## Results

### Single-cell profiles of several histone modifications in the mouse brain

To investigate the epigenetic landscape of OLGs in the juvenile brain, we coupled antibody directed tagmentation (Cut&Tag)^1^ with an existing single-cell ATAC-seq protocol (Figure 1a). For this purpose, several modifications to the Cut&Tag protocol were implemented, including optimization of bulk tagmentation of small number of nuclei extracted from primary sorted cells from the CNS (see Methods). For instance, to reduce clumping of the nuclei during the procedure, we included 1% BSA in specific buffers throughout the protocol (Extended Data Figure 1a). Addition of BSA reduced clumping of the nuclei (Extended Data Figure 1b) but did not significantly alter the efficiency of tagmentation nor the signal distribution for bulk Cut&Tag (Extended Data Figure 1c).

**Figure 1.**
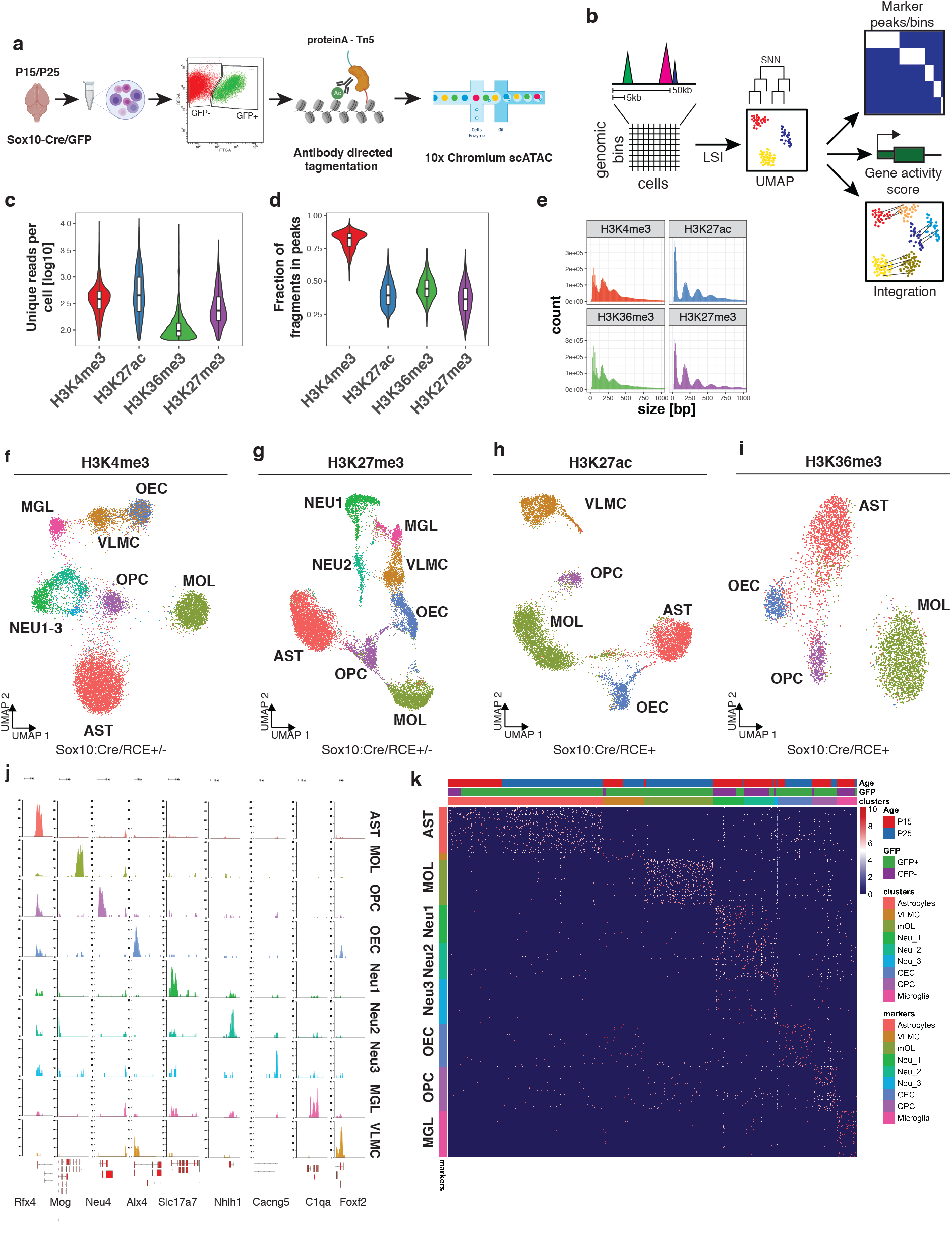
Single-cell profiling of several histone modifications in the mouse brain. a. Schematic of the scCut&Tag experimental design. Cells were isolated from mouse brain at age P15 or P25, sorted either for GFP+ or GFP-population; nuclei were isolated and incubated with specific antibody against chromatin modifications or transcription factors and tagmented using proteinA-Tn5 fusion and processed by the 10x chromium scATAC-seq protocol. b. Schematic of the analysis strategy. scC&T signal was aggregated into cell x bin matrix, with various bin size (5kb or 50kb); dimensionality reduction was performed using LSI and UMAP and clustering using SNN. Cell clusters were used to identify marker regions, gene activity scores was calculated per cell and integration with other datasets was performed. c. Comparison of number of unique reads per antibody in scC&T experiments. d. Comparison of percentage of reads falling into peak regions per antibody in scC&T experiments. Peaks were obtained by peak calling in merged bulk datasets. e. Distribution of fragment lengths scC&T experiments per antibody. f-i. two-dimensional UMAP representation of the scC&T data for H3K4me3, n=4 (f), H3K27me3, n=4 (g), H3K27ac, n=2 (h) and H3K36me3, n=2 (i). AST – Astrocytes, MOL – Mature Oligodendrocytes, OPC - Oligodendrocyte progenitor cells, NEU – Neurons, MGL - Microglia, VLMC – Vascular and leptomeningeal cells, OEC – Olfactory ensheating cells. j. Pseudo-bulk scC&T profiles of H3K4me3 aggregated by cell type at markers’ loci. k. Heatmap showing H3K4me3 signal intensity in top 50 most specifically enriched genomic bins per cluster (rows) and single cells (columns). Cells are randomly sampled and 5% of total number of cells are displayed. Color bars in rows specify marker cluster, and in columns cell metadata (Age, GFP).

We aimed to use scC&T to investigate epigenetic changes during OLG differentiation and myelination in the mouse brain. We used a mouse model expressing Sox10:Cre/Rosa26:(CAG-LSL-EGFP) mice (RCE)^12,16,17^, which primarily labels OLGs in the mouse CNS. We sorted both GFP+ and GFP-cells from whole juvenile brain at postnatal day 15 (P15 timepoint). Additionally, to gain more insights into OLG differentiation, we sorted GFP+ cells at P21-P25 range (P25 timepoint) in two biological replicates, at the peak of oligodendrocyte differentiation and the onset of myelination (Figure 1a). We then performed scC&T with antibodies against H3K4me3 (for active promoters) and H3K27me3 (for repressed loci) at both timepoints. We also performed scC&T against H3K27ac (marks active enhancers and promoters) and H3K36me3 (marks actively transcribed genes) in the P25 GFP+ cohort. We obtained between median of 98 (H3K36me3) and 453 (H3K27ac) unique fragments per cell (Figure 1c) of which between 39.4% to 85.6% of fragments fell within peak regions (Figure 1d), indicating a low level of background. The fragment length distribution was consistent with the capture of sub nucleosome fragments as well as mono-, di- and tri -nucleosomes for all modifications (Figure 1e).

Inspection of merged pseudo-bulk datasets including all cells revealed distinct profiles for the histone modifications highlighting specificity of the technique (Extended Data Figure 2a). In particular, H3K4me3 was mainly present at regions flanking transcription start sites, H3K27ac occupied both these regions and neighboring intergenic regions (most likely enhancers) and H3K36me3 was spread through the coding regions (Extended Data Figure 2a). In contrast, H3K27me3 was associated with genes where the other active markers were absent, or it coincided with H3K4me3. We identified single cells based on number of reads per barcode and fraction of reads falling into peak regions, called from merged bulk data (Extended Data Figure 2b and Methods). Altogether we obtained Cut&Tag profiles of various histone modifications for 47 340 single cells.

### scC&T of individual histone modifications allows identification of specific cell populations from the mouse brain

*De novo* identification of cell populations is one of the main strengths of a single-cell resolution techniques. To pursue this objective, we aggregated the data and generated cell-feature matrices for all datasets using 5kb (H3K4me3, H3K27ac and H3K27me3) or 50kb genomic bins (H3K36me3) (Figure 1b). To reduce the dimensionality of the dataset, we used latent semantic indexing (LSI) and uniform manifold approximation and projection (UMAP) and clustered the cells using shared nearest neighbors (SNN) and Leiden algorithm implemented with package Signac/Seurat v3 (Figure 1b).

We found all major CNS cell populations in the scC&T assay for the different modifications (Figure 1f-i), by identifying specific peaks proximal to promoters of marker genes (Figure 1j,k, Figure 2). We manually annotated the populations as mature oligodendrocytes (MOL, *Mbp*+, *Mog*+, *Cldn11*+), Astrocytes (AST, *Slc1a2*+, *Rfx4*+, *Aqp4*+), olfactory ensheathing cells (OEC, *Alx3*+, *Alx4*+, *Frzb*+), vascular cells (VAS, *Nes*+, *Tbx18*+, *Foxf2*+) and cells within the spectrum of oligodendrocyte progenitors cells (OPCs), committed OPCs (COPs) and newly formed oligodendrocytes (NFOLs) (OPC, *Pdgfra*+, *Neu4*+, *Gpr17*+) in the GFP+ fraction (Figures 1j, 2, Extended data figure 3). GFP-cells were comprised mainly of Neurons (NEU, *Rbfox3+, Neurod2+*), both Excitatory (Exc, *Slc17a7+*) and Inhibitory (Inh, *Gad1+, Gad2+*), Astrocytes (AST, *Slc1a2*^+^, *Rfx4*^+^, *Aqp4*^+^) and Microglia (MGL, *C1qa+, CD45/Ptprc+*) (Figures 1j, 2, Extended data figure 3). We could identify similar populations in H3K27me3 scC&T and annotate them using a combination of markers that lacked the repressive mark in the vicinity of the marker gene regions (Figure 1j, 2, Extended data figure 3).

**Figure 2.**
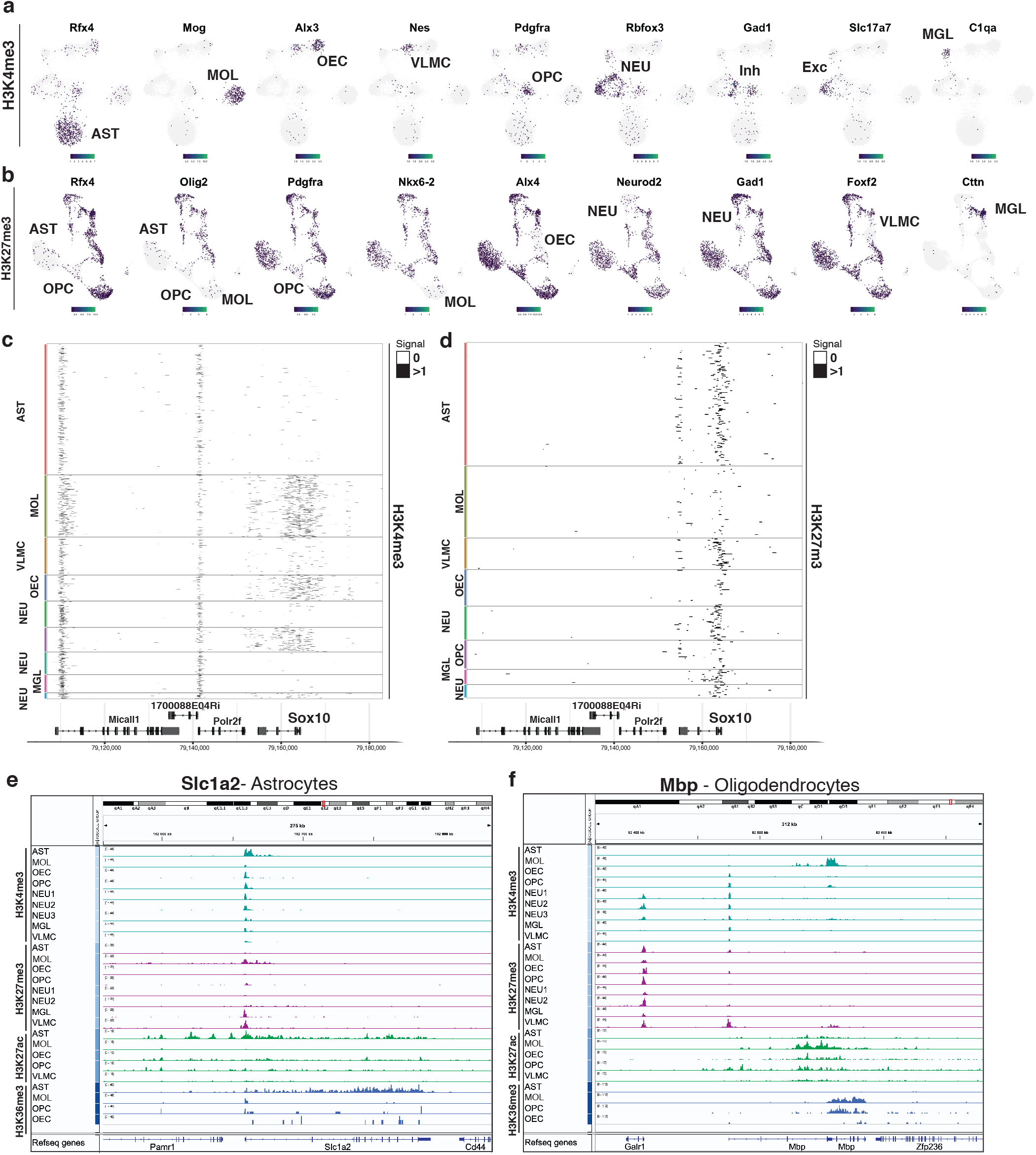
De novo identification of cell types by cell type specific scCut&Tag marker regions. a-b. Projection of scCut&Tag gene activity scores of (a) H3K4me3 and (b) H3K27me3 on two-dimensional UMAP embedding. Gene name is depicted in the title and specific population is highlighted in the UMAP plot by labeling the cell type. c-d. Heatmap representation of the scC&T signal for (c) H3K4me3 and (d) H3K27me3. X axis represents genomic region, each row in Y axis contains data from one cell. Cell correspondence to clusters is depicted by color bar on the right side of the heatmap and annotated with cell type. Signal is aggregated per 250 bp windows and binarized. e-f. Aggregated pseudobulk scC&T profiles for four histone modifications in all identified cell types at the loci of selected marker genes (Slc1a2 – astrocytes and Mbp – oligodendrocytes)..

The clustering was highly reproducible among biological replicates and cells originating from P15/P25 age were well intermingled within the clusters (Extended Data Figure 2d-f). The cell states of OLG lineage reflect the mouse age with majority of OPCs originating from P15 and differentiated OLG coming from P25 (Extended Data Figure 2d, 2f). Interestingly, we detected a major, likely transient population of astrocytes, in GFP+ fraction of the Sox10:Cre/RCE mouse (Extended data Figure 2d, 2f), most probably derived from ventral regions^18–20^.

The scC&T data also allowed to discriminate distinct histone modification patterns between the diverse cell populations that are confounded in bulk approaches. As an example, the TSS of Sox10 appeared to be bivalent for H3K4me3 and H3K27me3 in the pseudo-bulk brain dataset (Extended Data Figure 2a), but single cell analysis indicated that only oligodendrocytes, OEC and OPC populations had active marks (H3K4me3, H3K27ac) deposited on the promoter of Sox10 (which was used to sort GFP+ cells), while astrocytes and vascular cells did not (Figure 2c,d). Occupancy of H3K36me3 was similarly found across the gene body of Sox10 in respective clusters, whereas the repressive mark H3K27me3 was specifically depleted from Sox10 promoter in oligodendrocytes, OPCs and OECs (Figure 2c,d, Extended Data Figure 3c,d).

### Integration of scC&T data with single-cell gene expression

In order to validate the manual annotation of the clusters, we used the adolescent mouse brain scRNAseq atlas^3^. We picked the one hundred most specifically expressed marker genes for selected populations, generated metagene modules and then calculated gene activity score (scC&T signal in gene body and promoter) within the module. We found that the specific cell clusters showed enrichment of the metagene signal for the active modifications and were depleted of the signal in the H3K27me3 dataset (Extended Data figure 4), supporting the cluster annotations. Furthermore, we integrated the H3K4me3 scCut&Tag with the scRNA-seq data using canonical correlation analysis (CCA)^21^ at the single cell level. We found that the major cell populations co-cluster together with the corresponding scRNA-seq population (Figure 3a). Finally, we used gene ontology (GO) terms analysis to functionally annotate the H3K4me3 scC&T clusters and found GO terms such as Astrocyte differentiation and activation (AST), Myelination (OLG), regulation of myelination (OEC), cell migration involved in vasculogenesis (VAS), glial cell development (OPCs), neuron development, neuron maturation and axonogenesis (NEU) and microglial cell activation involved in immune response (MGL) (Extended Data Figure 5) to be specifically enriched in respective clusters.

**Figure 3.**
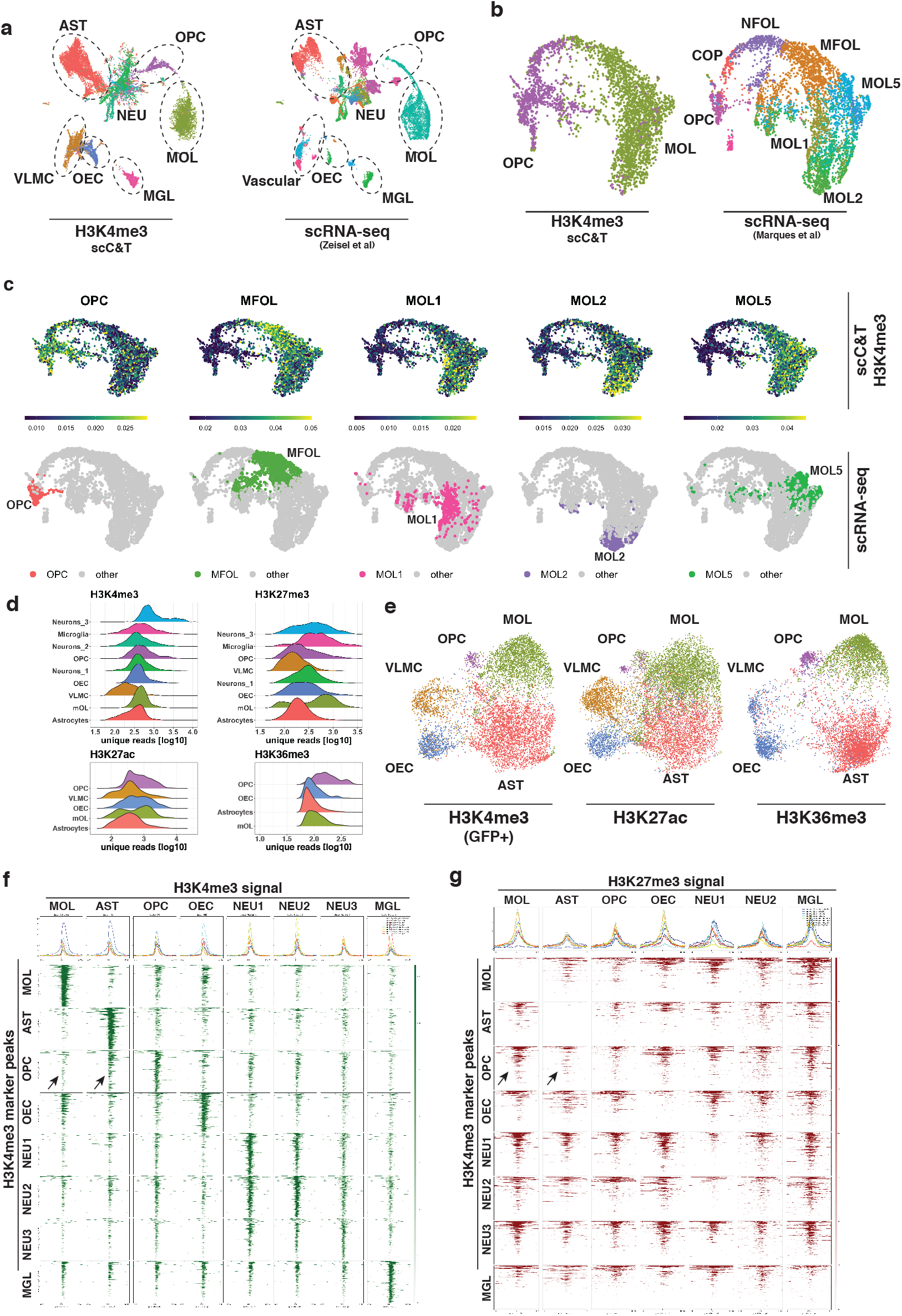
Integration of the scC&T data with single-cell gene expression. a. Coembedding of the H3K4me3 scC&T data together with the mouse brain atlas scRNA-seq data3. b. Coembedding of the H3K4me3 of OPC and MOL clusters together with the scRNA-seq data15 depicting heterogeneity of the OLG lineage. c. Metagene activity scores of OLG lineage sub-clusters. Activity scores are calculated by aggregating the H3K4me3 scC&T signal (in gene body and promoter) over the 200 most specific genes (determined by p-value) expressed in the corresponding scRNA-seq cluster15. The corresponding scRNA-seq populations are highlighted in the UMAP plots in the bottom row d. Ridgeline plots depicting histogram of number of unique reads per population of four histone modifications. e. Coembedding of the scC&T data of three active histone modifications – H3K4me3, H3K27ac and H3K36me3 in a single two-dimensional UMAP space. H3K4me3 dataset was filtered for cells corresponding to populations present in the GFP+ samples. f-g. Metagene heatmaps of (f) H3K4me3 and (g) H3K27me3 occupancy in populations (x axis) at the loci of H3K4me3 peaks (y axis). Arrows highlight the H3K4me3 and H3K27me3 signal in MOL and AST populations around the OPC marker regions. AST – Astrocytes, MOL – Mature Oligodendrocytes, OPC - Oligodendrocyte progenitor cells, NFOL – Newly formed oligodendrocytes, NEU – Neurons, MGL - Microglia, VLMC – Vascular and leptomeningeal cells, OEC – Olfactory ensheating cells.

The scC&T data was able to resolve the major cell types in a heterogenous sample. However, a previous study had shown that further cell subtypes can be detected in the population of oligodendrocytes^15^. Therefore, we asked whether this heterogeneity can be resolved by integration of the scC&T data with an existing OLG scRNA-seq dataset^15^. For this purpose, we co-embedded the H3K4me3 scC&T and scRNA-seq using CCA, which showed good integration of the techniques, while the scRNA-seq clustering was retained (Figure 3b). We then used metagene scores of OPC, MFOL, MOL1, MOL2 and MOL5 to reveal cell subtype signatures within the H3K4me3 scC&T data. Interestingly, we found that the population of oligodendrocytes that appeared homogenous could be further deconvoluted into sub-populations enriched in module-specific genes (Figure 3c), indicating that oligodendrocyte heterogeneity is reflected at an epigenetic level.

### Differential global and genome-wide patterns of histone modifications in the single-cell populations

Since the scC&T profiles are generated simultaneously for all present populations, it allows for quantitative analysis of global and genome-wide pattern of histone modifications in them. We used the number of unique reads per cell as a proxy for absolute amount of the histone modification in single cells. We found a substantial variability in this regard (Figure 3d). This is most prominent for H3K27me3, which is enriched in populations of oligodendrocytes, microglia and a subset of neurons, relative to the other populations (Figure 3d). Interestingly, we also observed relatively higher amounts of H3K36me3 in the population of immature oligodendrocytes (OPC/COP-NFOL stages) (Figure 3d).

Next, we asked whether we could assign cell populations across the different active modifications and cross-correlate them. For this purpose, we used the CCA to integrate the data at gene resolution. Strikingly, the obtained two-dimensional representation of the data recapitulated the original non-supervised clustering with great precision (Figure 3e) and the clusters annotated with the same cell type in different datasets co-occupied the same low dimensional space (Figure 3e). To further look at the interplay between active and repressive marks, we identified all active promoters specific for individual populations marked by H3K4me3 and plotted signal of H3K4me3 (Figure 3f) and H3K27me3 (Figure 3g) per cluster for all the populations. As expected, we observed depletion of H3K27me3 when the promoter is enriched in H3K4me3 in the respective population (Figure 3g). Interestingly, we noticed that there was low occupancy of H3K4me3 marks on OPC genes in differentiated OLG, whereas these genes had higher H3K4me3 signal in astrocytes (Figure 3f). Besides, H3K27me3 signal was depleted from OPC-specific genes in astrocytes (Figure 3g), suggesting that H3K27me3 is not required to repress OPC genes during astrocyte differentiation. Consistently, it was recently reported that disruption of H3K27me3 impairs OPC differentiation to MOL, and triggers switch towards an astrocytic faith^22^. Moreover, the presented epigenetic profiles suggest that astrocytes are epigenetically (H3K4me3 and H3K27me3) more similar to OPCs than MOLs are.

### Increase in breadth of H3K4me3 upon oligodendrocyte differentiation

The breadth of the H3K4me3 mark has been previously linked to cell identity, gene expression and transcriptional consistency across a variety of cell types^23^. We noticed in the H3K4me3 pileup analysis that both amplitude and breadth of the H3K4me3 signal were increased at population-specific marker gene promoters, when compared to marker gene promoters of other populations (Extended Data figure 3f). To quantify the breadth, we specifically looked at promoters of marker genes identified from scRNA-seq data. We found that the marker genes of the identified populations had on average higher H3K4me3 breadth (Figure 4a). Moreover, the magnitude of the breadth was different for individual cell types with the broadest H3K4me3 peaks on the promoters of astrocytes and oligodendrocytes, and the narrower on VLMCs, but also OPCs (Figure 4b), which could suggest an increase of H3K4me3 breadth during transition from progenitor states to fully differentiated states. To further look into the dynamics of H3K4me3 spreading, we leveraged the ability of scCut&Tag to visualize H3K4me3 spreading at the single cell resolution and investigated the H3K4me3 breadth during the process of differentiation of MOLs from OPCs. We generated single-cell H3K4me3 metagene profiles around MOL specific marker genes. We then ordered the cells (OPCs and MOL) in the matrix according to H3K4me3 signal coming from genes expressed in MOLs (MOL signature) and validated by pseudotime analysis (Figure 4c, 4d). Strikingly, we observed a gradual increase in the breadth of the H3K4me3 signal at MOL promoters with single cell resolution (Figure 4e), which is consistent with spreading of H3K4me3 as the cells progress towards differentiated oligodendrocyte identity.

**Figure 4.**
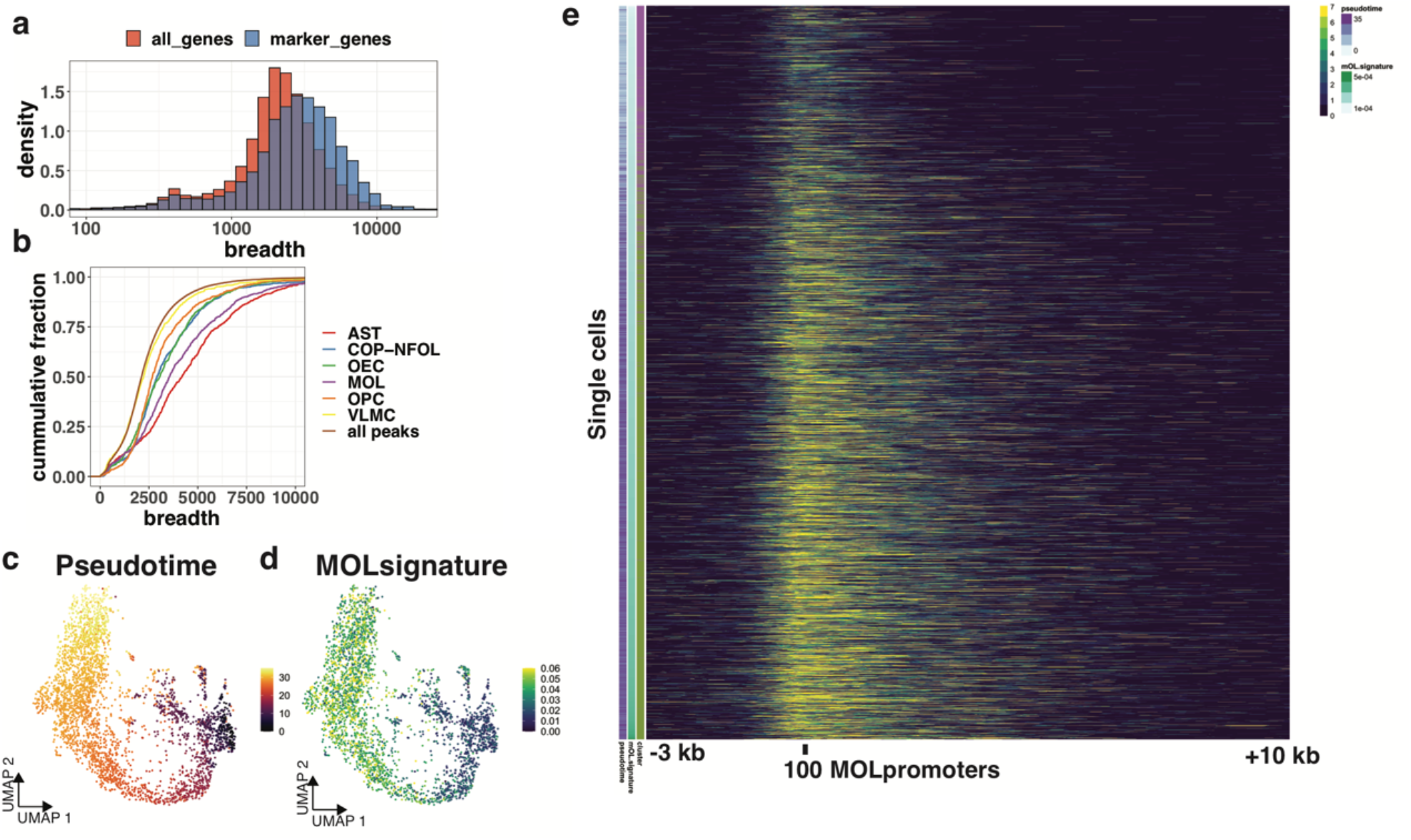
Spreading of H3K4me3 mark from promoters at single-cell resolution. a. Histogram of breadth of H3K4me3 around promoters of all annotated genes (all_genes) and genes that are marker genes for specific populations in scRNA-seq (marker_genes). b. Cumulative distribution of the breadth of H3K4me3 peaks around the promoters of marker genes identified from scRNA-seq. c. Pseudotime analysis of the OLG lineage H3K4me3 scC&T data that was integrated with scRNA-seq data15. d. MOL signature score (H3K4me3 signal in MOL-specific gene body and promoter normalized to total number of reads in each cell) projected on the UMAP representation of H3K4me3 scC&T data. e. Heatmap depicting H3K4me3 spreading from the promoters of MOL-specific genes. Each row represents a single cell. Cells are ordered by the MOL signature score, that is correlated also with pseudotime. X axis shows the genomic distance from the meta-promoter (−3kb / +10kb). Meta-promoters are promoters of top 100 most specifically expressed genes in MOLs defined by scRNA-seq data.

### Single cell Cut&Tag of transcription factors

Transcription factor binding is notoriously difficult to profile using ChIP-seq in low input samples. Therefore, we asked whether scC&T was able to uncover the binding of transcription factors at a single-cell resolution. We chose the transcription factor (TF) Olig2 since it is specific for glial populations, and Rad21, a general chromatin architecture factor and a subunit of the cohesin complex. We performed scC&T in GFP+ sorted cells from the brain at postnatal day P25. The number of unique reads per cells was lower for TF scC&T compared to histone modifications. Nevertheless, we were able to obtain in median 48 and 240 unique reads per cell in Olig2 and Rad21 scC&T respectively. We reduced the dimensionality of the dataset using LSI and UMAP and obtained specific clusters for Rad21 and Olig2 (Figure 5a-d). Olig2 dataset separated into 2 clusters based on depth (number of unique reads per cell), and we annotated the cluster with low number of unique reads as “low binders” (Figure 5a). Since manual annotation of populations based on markers is challenging in TF Cut&Tag (Figure 5e), we based cluster annotation on the assumption that Olig2/Rad21 binding in specific cell types is correlated with enhancer/promoter activity. Therefore, we analyzed Olig2/Rad21 binding in promoter regions of genes that are specifically modified by H3K4me3 in scC&T data, and identified populations of AST, OLG and OEC in Rad21 scC&T and OLG and non-OLGs (‘low binders’) in Olig2 scC&T (Extended data figure 6a-d). The OLG population in Olig2 scC&T is likely composed of both mature OLG and OPCs, which appear to form a subpopulation within the OLG cluster (Extended data figure 6a). To further strengthen the cluster annotations, we performed co-embedding of the Rad21/Olig2 with another histone modification – H3K27ac using CCA and found that the identified clusters consistently co-embeded with the corresponding H3K27ac clusters (Extended data figure 6e). Interestingly, the low binder non-OLGs cells randomly co-embedded with the populations of OEC, Astrocytes and vascular cell clusters that were defined by H3K27ac, whereas OLG cluster specifically co-embeded with the OLG H3K27ac signal (Extended data figure 6e). This finding is consistent with expression of Olig2 throughout the cell types, being highly expressed throughout the OLG lineage, whereas OECs and VLMCs do not express Olig2 mRNA and only small portion of astrocytes is expressing Olig2 (Figure 5f).

**Figure 5.**
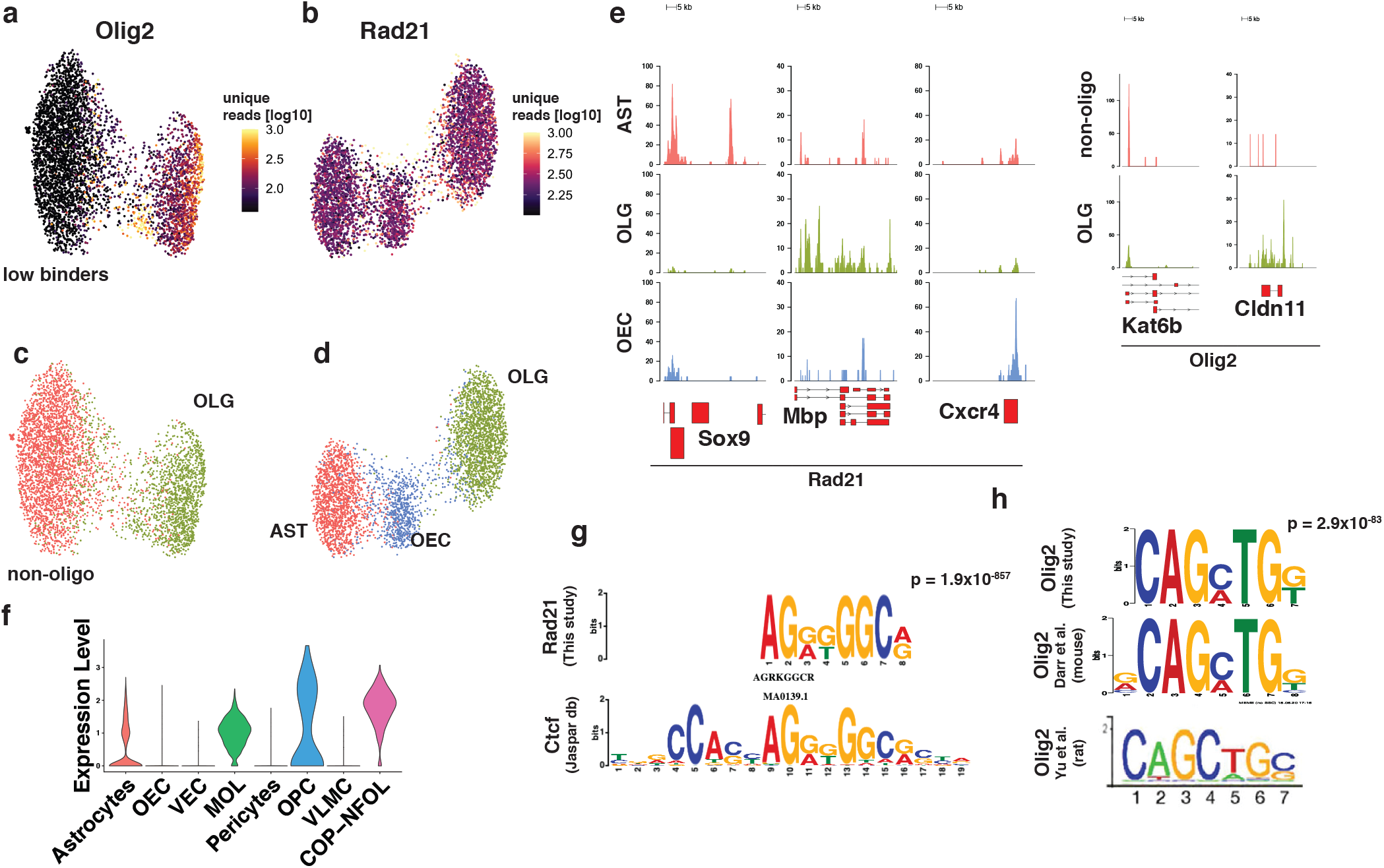
scC&T analysis of transcription factors binding. a-b. Two dimensional UMAP embedding of (a) Olig2 and (b) Rad21 scC&T data with coloring by the number of unique reads per cell. c-d UMAP embedding of the (c) Olig2 and (d) Rad21 scC&T with coloring by the cell type. e. Pseudobulk profiles of Olig2 and Rad21 scC&T data aggregated by cell type around the marker gene regions. f. RNA expression of Olig2 from scRNA-seq data for cell types present in Sox-10-Cre/RCE+ populations. g. Logo representation of the motif that was found to be the most enriched in the scC&T of Rad21 and its alignment with the motif of transcription factor Ctcf retrieved from the Jaspar database. h. Logo representation of the motif found to be enriched in the Olig2 Cut&Tag that is consistent with the previously reported Olig2-specific motif26,27. AST – Astrocytes, MOL – Mature Oligodendrocytes, OPC - Oligodendrocyte progenitor cells, NEU – Neurons, MGL - Microglia, VLMC – Vascular and leptomeningeal cells, OEC – Olfactory ensheating cells.

To validate the specificity of the scC&T, we searched for enriched motifs in binding sites using MEME suite^24^ in merged pseudo-bulk datasets of Rad21 and Olig2. We found the motif of chromatin architecture factor, CTCF as the highest enriched in the Rad21 dataset (Figure 5g), which is consistent with the cooperativeness between Ctcf and Cohesin^25^. We found several motifs enriched in the Olig2 scC&T, including motif CAGMTG, similar to the previously identified CAGMTG/CAGCTG motif specific for Olig2 (Figure 5h) in mouse^26^ and rat^27^ respectively. Together with the previously identified Olig2 motif, we found enrichment of multiple general eukaryotic enhancer and promoter features (GC box, CAAT box). Interestingly, we also found a motif similar to the motif of transcription factors from the Sox family (ACARWR, Extended Data Figure 6f), which is consistent with physical interaction and cooperativity between Olig2 and members of Sox family of transcription factors (Sox8, Sox10)^28^.

### Prediction of enhancer-promoter interactions from scCut&Tag data

Perhaps the most intriguing and challenging task of epigenomics is to use epigenetic data to predict gene regulatory networks. Our scC&T dataset is a rich resource that can be used to tackle this question in great detail. We used the activity-by-contact model (ABC model)^29^ of enhancer-promoter interactions to predict the gene-enhancer regulatory networks from aggregated scC&T data (Figure 6a). We focused on OLG and ran the ABC model using oligodendrocyte H3K27ac scC&T, oligodendrocyte scATAC-seq data (P50 cortex, 10x genomics) and Hi-C of neural progenitor cells^30^. By using the neural progenitor cells Hi-C we ensured that we did not provide the model with accurate measurements of DNA looping, but rather used the Hi-C data to estimate the long-distance chromosome topology. The ABC model predicted ∼200,000 enhancer-promoter loops in the oligodendrocyte lineage. In order to validate the predictions, we performed HiChIP of H3K27ac from primary mouse OPC cultures that are known to contain OPCs and differentiated oligodendrocytes. To examine the specificity of the predicted loops, we generated pileups of Oligodendrocyte H3K27ac HiChIP data and H3K27ac HiChIP performed on mouse embryonic stem cells^31^ and found that the predicted loops were highly specific for oligodendrocytes (Figure 6b). We then used the scC&T data to further refine the loops. We used the oligodendrocyte population H3K4me3 signal to filter for promoters that are active in oligodendrocytes, which yielded 61,000 loops. Moreover, we presumed that cohesin (Rad21) binding is correlated with probability of loop formation and filtered interactions that have Rad21 binding on both sides of the loop (promoter and enhancer), resulting in ∼5,000 loops. These high confidence loops show significantly higher specificity in the HiChIP pileup signal, compared to the original set of unfiltered loops (Figure 6c). Moreover, loops predicted by ABC model and filtered by H3K4me3 signal overlapped well with H3K27ac-mediated loops and stripes, when inspected in the 2D matrix representation (Figure 6d).

**Figure 6.**
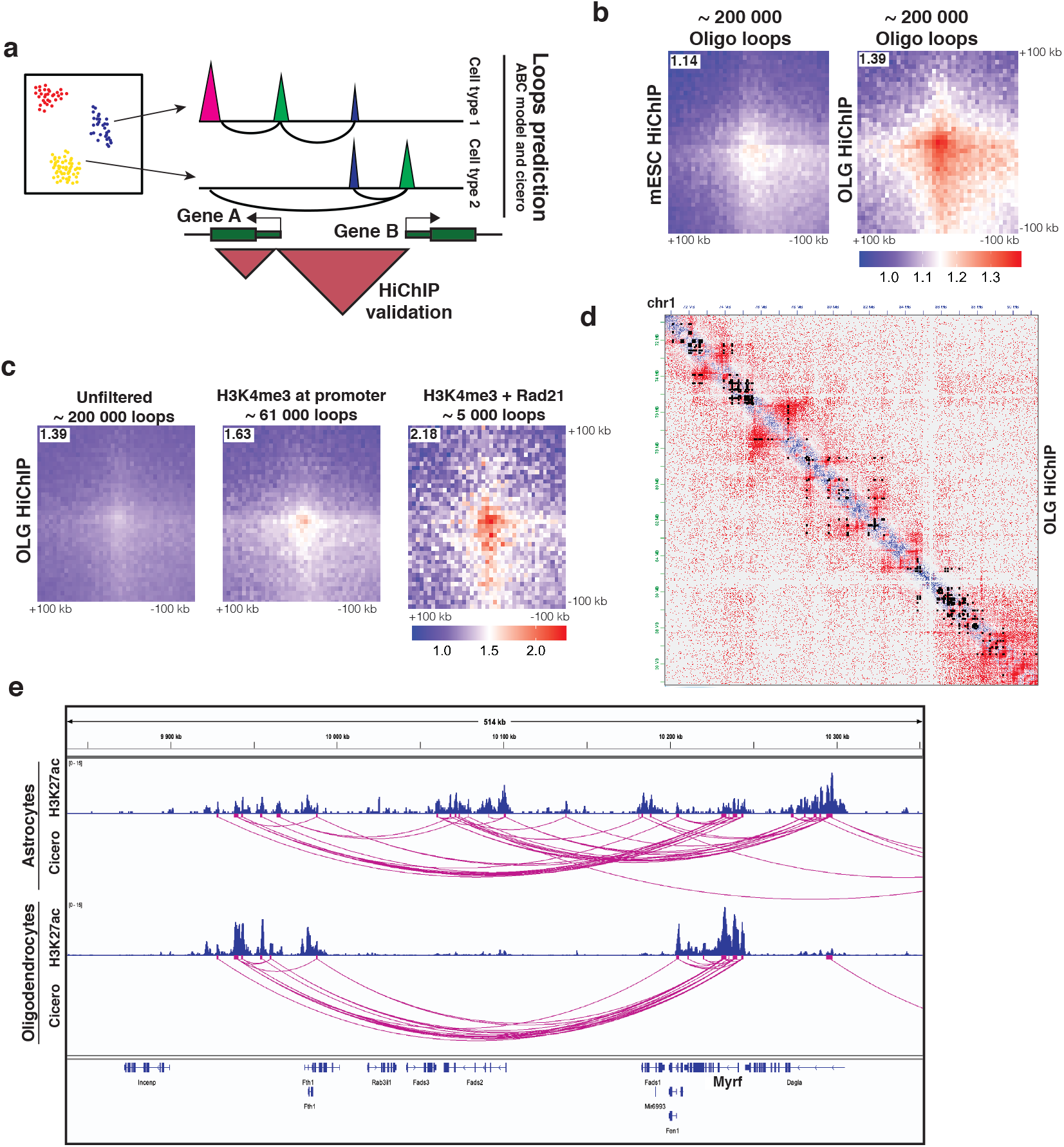
Prediction of gene regulatory networks from the scC&T data. a. Schematic depicting the strategy used to predict and validate the promoter-enhancer specific loops. Loops are predicted using the H3K27ac scC&T and other publicly available datasets (see Methods). Presence of loops is validated by H3K27ac HiChIP analysis of purified populations of OLG lineage b. Pileup analysis of 200,000 loops predicted by the ABC model. Signal of HiChIP performed in either mES cells or OLG lineage cells was aggregated and plotted as a heatmap with the center at the intersection of the loop coordinates. c. Pileup analysis of loops predicted by ABC and filtered using H3K4me3 (∼61,000 loops) and Rad21 (∼5000 loops) scC&T data. d. Representative overlay of OLG HiChIP matrix with the loops predicted by the ABC model from the scC&T data and filtered with the H3K4me3 signal (black dots). e. Representative example of the loops predicted with cicero specific for MOL and AST populations. Cicero using H3K27ac scC&T predicts a regulatory region ∼300kb upstream of the Myrf gene specific for MOL.

Finally, cicero^32^ can be used to predict cell type specific interactions of cis-regulatory regions through a different approach - correlation of ATAC-Seq signal in single cells. Applying cicero to the scC&T data, we found 21,626 H3K27ac features with correlation score > 0.2. As an example, we extracted correlated features from the surroundings of an important oligodendrocyte-specific transcription factor Myrf (Figure 6e) for which we predicted a distal regulatory element ∼300 kb upstream of the Myrf gene. This element was specific for oligodendrocytes with different topology in Astrocytes, consistent on the specific expression of Myrf in the former population. Altogether, predictions of distal regulatory elements based on scC&T data can serve as an important tool to guide future studies and/or perturbations of cell type specific regulatory elements.

## Discussion

The emergence of single-cell resolution transcriptomic technologies has allowed the investigation of developmental biology processes with exquisite detail. However, the underlying regulatory epigenetic processes still remain to be uncovered at such resolution.

Here, we provide for the first time a high-throughput study of chromatin modifications and transcription factor binding at single-cell resolution in the mouse brain. We generated an extensive dataset of histone modifications in juvenile mouse at the age of P15-P25, at the time of differentiation of OPCs into mature oligodendrocytes and at the peak of myelination. We generated H3K4me3 and H3K27me3 genome tracks for all major cell types in the brain, while providing H3K27ac and H3K36me3 tracks for glial cell populations. We show that analysis of histone modifications at a single-cell level allows identification of distinct neural cell types, with the active marks H3K4me3 and H3K27ac, and the repressive mark H3k27me3 being particularly powerful in this aspect.

The main strength of scC&T when applied to tissues lies in its unbiased and unsupervised nature. We were able to *de novo* identify distinct populations of cells and find unique peaks of histone modifications at genomic elements such as promoters, enhancers and gene bodies of marker genes. This allows the examination of dynamics of epigenetic changes in virtually any biological process including in development and disease, without *a priori* knowledge about the process. This attribute makes scC&T an extremely valuable tool that can be used to study epigenetic regulation of gene expression. Moreover, with low background signal, scC&T can yield merged genomic tracks from as little as 500 cells with quality comparable to bulk ChIP-seq performed on hundreds of thousands to millions of cells.

We further used the histone modifications scC&T data to examine epigenetic concepts such as promoter bivalency, spreading of H3K4me3 from the promoters and enhancer-promoter connectivity. scC&T allowed deconvolution of potential H3K4me3/H3K27me3 bivalency, but also a close epigenetic relationship between OPCs and Astrocytes regarding H3K4me3 and H3K27me3. We also observed an increase in H3K4me3 breadth during the process of differentiation in the OLG lineage, indicating that spreading of this modification can be used to delineate lineage progression. We also show that various histone modifications and transcription factor binding can be incorporated into models of gene regulatory networks in primary cell types at various developmental timepoints at high throughput. Acquiring such detailed data was previously possible only for cultured cell lines, at whole tissue resolution^33^, or after detailed characterization of cell surface markers that could be used for sorted cell populations.

We also generated single cell scC&T profiles for glial transcription factor Olig2 and chromatin architecture factor Rad21. While Olig2 preferentially binds oligodendrocyte lineage specific enhancers and promoters and binds only weakly to astrocyte specific gene regulatory regions, Rad21, a ubiquitously expressed genome architecture factor, binds cell type specific promoters and enhancers. We were able to distinguish cell identity based only on Rad21 binding for all cell types, except for vascular cells and OPCs, which are the smallest cell populations and share features with larger populations. The Rad21 binding at single-cell resolution provides valuable insights into enhancer-promoter connectivity and we show that it can be incorporated into models that aim to uncover enhancer-promoter regulatory networks.

Although the provided datasets provide unique insights into epigenetic regulation, improvement of the technology, especially number of unique fragments per cell can further enhance the possible applications. In the current state, scC&T is able to distinguish the major cell types but fails *per se* to uncover heterogeneity of the subpopulations in an unsupervised manner. We could only reveal the oligodendrocyte heterogeneity at H3K4me3 level upon integration with more complex scRNA-seq data. Potential introduction of multi-omic approaches, such as simultaneous measurement of scC&T and RNA-seq signal from the same cell will help us to further understand causal relationships between epigenetic modifications and gene expression.

## Methods

### Animals

The mouse line used in this study was generated by crossing Sox10:Cre animals^7^ (The Jackson Laboratory mouse strain 025807) on a C57BL/6j genetic background with RCE:loxP (EGFP) animals^8^ (The Jackson Laboratory mouse strain 32037-JAX) on a C57BL/6xCD1 mixed genetic background. Females with a hemizygous Cre allele were mated with males lacking the Cre allele, while the reporter allele was kept in hemizygosity or homozygosity in both females and males. In the resulting Sox10:Cre-RCE:LoxP (EGFP) animals the entire OL lineage was labeled with EGFP. Breeding with males containing a hemizygous Cre allele in combination with the reporter allele to non-Cre carrier females resulted in offspring where all cells were labeled with EGFP and was therefore avoided. All animals were free from mouse viral pathogens, ectoparasites and endoparasites and mouse bacteria pathogens. Mice were kept with the following light/dark cycle: dawn 6:00-7:00, daylight 7:00-18:00, dusk 18:00-19:00, night 19:00-6:00 and housed to a maximum number of 5 per cage in individually ventilated cages (IVC sealsafe GM500, tecniplast). Cages contained hardwood bedding (TAPVEI, Estonia), nesting material, shredded paper, gnawing sticks and card box shelter (Scanbur). The mice received regular chew diet (either R70 diet or R34, Lantmännen Lantbruk, Sweden). Water was provided by using a water bottle, which was changed weekly. Cages were changed every other week. All cage changes were done in a laminar air-flow cabinet. Facility personnel wore dedicated scrubs, socks and shoes. Respiratory masks were used when working outside of the laminar air-flow cabinet. Animals were sacrificed at juvenile stages (P21-25) and both sexes were included in the study. All experimental procedures on animals were performed following the European directive 2010/63/EU, local Swedish directive L150/SJVFS/2019:9, Saknr L150 and Karolinska Institutet complementary guidelines for procurement and use of laboratory animals, Dnr. 1937/03-640. The procedures described were approved by the local committee for ethical experiments on laboratory animals in Sweden (Stockholms Norra Djurförsöksetiska nämnd), lic. nr. 131/15, 144/16 and 1995_2019.

### List of antibodies

**Table 1.** List of used antibodies.

| Antibody | type | Manufacturer | Product no. |
| --- | --- | --- | --- |
| H3K4me3 | primary | Diagenode | C15410030 |
| H3K27ac | Primary | Abcam | Ab177178 |
| H3K27me3 | Primary | Cell signaling | 9733T |
| H3K36me3 | Primary | Abcam | Ab9050 |
| Rad21 | Primary | GeneTex | GTX106012 |
| Olig2 | primary | Novus Biologicals | NBP1-28667 |
| Guinea pig anti-rabbit | Secondary | Novus Biologicals | NBP1-72763 |

### Single cell Cut&Tag

Single cell Cut&Tag was performed as in Kaya-Okur *et* al.^2^ with modifications. The Cut&Tag was performed in 0.5 ml tubes, all washes and incubation volumes were 200 ul unless otherwise stated and all centrifugations were done using swinging bucket centrifuge, with appropriate tube adaptors.

150,000 GFP+ or GFP-cells were sorted from the brain of Sox10::Cre/RCE animals on post-natal day 15 (P15) or 21-25 (P21-P25) using fluorescent assisted cell sorting (FACS Aria III or FACS Aria Fusion) directly into the Antibody buffer (20 mM HEPES pH 7.6, 150 mM NaCl, 2mM EDTA, 0.5 mM Spermidine, 0.05% Digitonin, 0.01 % NP-40, 1x Protease inhibitors, 1% BSA), centrifuged for 5 minutes @300 x g force, washed 1x with the antibody buffer, resuspended in 200 ul antibody buffer with 1:50 diluted primary antibody (Table 1) and incubated overnight with slow rotation. Next day the nuclei were centrifuged 3 minutes @ 600 x g, washed 1x with 200 ul of Dig-wash buffer (20 mM HEPES pH 7.6, 150 mM NaCl, 0.5 mM Spermidine, 0.05% Digitonin, 0.01 % NP-40, 1x Protease inhibitors, 1% BSA), resuspended in 200 ul of Dig-wash buffer with 1:50 diluted secondary antibody (Table 1) and incubated for 1h at room temperature with slow rotation. Then, nuclei were centrifuged 3 minutes @ 600x g, washed three times with 200 ul of Dig-wash buffer, resuspended in 200 ul of Dig-300 buffer (20 mM HEPES pH 7.6, 300 mM NaCl, 0.5 mM Spermidine, 0.05% Digitonin, 0.01 % NP-40, 1x Protease inhibitors, 1% BSA) with 1:100 diluted proteinA-Tn5 fusion and incubated for 1 hour rotating at room temperature. After Tn5 binding, the cells were centrifuged 3 minutes @ 300 x g and washed 3x with Dig-300 buffer, resuspended in 200 ul of tagmentation buffer (20 mM HEPES pH 7.6, 300 mM NaCl, 0.5 mM Spermidine, 0.05% Digitonin, 0.01 % NP-40, 1x Protease inhibitors, 10 mM MgCl^2^), without BSA and incubated for 1 hour at 37°C. After that, tagmentation was stopped by addition of 10 ul of 500 mM EDTA and 200 ul of Dig-300 buffer with BSA, and mixed by gently pipetting up and down several times (Final 0.5 % BSA, critical, otherwise the nuclei would clump during the centrifugation). Then the nuclei were centrifuged 3 minutes @ 300 x g, washed once with 1xPBS + 1% BSA and resuspended in 50 ul of 1x PBS+ 1% BSA before counting with counting chamber with trypan blue staining. At this stage clumping of the nuclei can be checked (with either trypan blue stain or DAPI) and if the sample is sufficient quality, processed using 10x chromium single cell ATAC-seq kit, skipping the Step 1 “Transposition” and continuing from Step 2.0 Generation & barcoding. We used version 1 chemistry of single cell ATAC-seq kit (10x genomics) according to the manufacturer’s instructions (https://assets.ctfassets.net/an68im79xiti/4dXLdjCzh5xKBlgRBkfSmW/eb0b5b463ced5778fa68e7dc50829e50/CG000168_ChromiumSingleCell_ATAC_ReagentKits_UserGuide_RevD.pdf)

### Single cell Cut&Tag data processing

Data was pre-processed using cell ranger-ATAC v1.2.0 (10x genomics), with standard parameters. However, cell identification was done manually using number of reads per cell and fraction of reads in peaks. Peaks were called using MACS^34^, single cell clustering and marker search was done using packages Seurat and Signac^21^, pseudotime analysis using slingshot^35^, motif search using MEME suite^24^, and compared to the Jaspar database^36^ or published ChIP-seq data, metagene plots using deepTools^37^ and custom scripts, most plots were generated using ggplot2 ^38^ R package, gene regulatory networks were analysed using ABC model^29^ and cicero^32^. Processing pipeline was build using Snakemake platoform^39^. Preprocessing pipeline and R notebooks used to generate the figures are shared at https://github.com/Castelo-Branco-lab/scCut-Tag_2020.

### proteinA-Tn5 production

The 3xFLAG pA-fusion sequence was copied from Addgene plasmid #124601 and inserted into psfTn5 plasmid (addgene #79107) to generate pA-Tn5 fusion construct (6xhis-TEV-3xFlag-pA-Tn5). The protein was purified from 3 litres of E.coli culture grown at in the LEX system, with protein expression temperature 18 degrees Celsius and cultivation temperature 30 degrees celsius. The temperature was switched to 18°C at OD=2 and expression induced at OD=3 with 0.5 mM IPTG. Bacteria were disrupted by sonication (4s/4s, 3min, 80% amplitude), centrifuged for 20 min at 49 000x g, filtered through 0.45 um filter and loaded on AKTA express column and purified overnight. 5 ml of HisTrap (GE Healthcare) was used for the affinity purification and Gel filtration was performed on HiLoad 16/60 Superdex 200 (GE Healthcare). Fractions were examined on SDS-PAGE gel before pooling, then desired fractions were combined, sample was diluted 1:5 to contain final 50% glycerol, aliquoted in 200 ul aliquotes and snap frozen using liquid nitrogen. Enzyme was stored at -80°C until loading.

### Tn5 loading

To begin with the complex formation, each of Mosaic end-adapter A (Tn5ME-A, TCGTCGGCAGCGTCAGATGTGTATAAGAGACAG) and Mosaic end-adapter B (Tn5ME-B, GTCTCGTGGGCTCGGAGATGTGTATAAGAGACAG) oligonucleotides were annealed with Mosaic-end reverse oligonucleotides (Tn5MErev, 5’-[phos]CTGTCTCTTATACACATCT-3’). To anneal, the oligonucleotides were diluted to 100µM. Each pair of oligos, Tn5MErev/Tn5ME-A and Tn5MErev/Tn5ME-B, was mixed separately resulting in 50µM annealed products respectively. The product was denatured in a thermocycler for 5 min at 95°C and then cooled down slowly in the thermocycler by turning off the thermocycler. The loading of pA-Tn5 with the oligonucleotides was carried out by incubating 2ul 50uM Tn5MErev/Tn5ME-A, 2ul 50uM Tn5MErev/Tn5ME-B, 21.56ul Glycerol, 21.3ul 2X Dialysis buffer (100 mM HEPES-KOH pH 7.2, 0.2 M NaCl(Thermo Fisher Scientific, AM9760G), 0.2 mM EDTA(Thermo Fisher Scientific, AM9260G), 2 mM DTT, 0.2% Triton X-100 (Thermo Fisher Scientific, 85111) and 20% Glycerol), 3.14ul Tn5 (63.64uM (3.5 mg/ml)), for 1h at room temperature (the final concentration is 2uM). The enzyme was stored at -20°C until further use, or at -80 for long term storage.

### Tissue dissociation

Mouse were sacrificed, perfused with 1xPBS and brain was removed. The brain was dissociated into single cell suspension using the Neural Tissue Dissociation Kit P (Miltenyi Biotec, 130-092-628) according to the manufacturers protocol. For mouse older than P7, myelin was removed using debris removal solution (Miltenyi Biotec, 130-109-398) according to manufacturer’s instruction. Single cell suspension was filtered through 50 um cell strainer and briefly stained with 1:5000 diluted DAPI (1mg/ml) to assess cell viability. For FACS, cells were resuspended in 1xPBS supplemented with 1% BSA and 2mM EDTA and kept at 4 degrees until sorted.

### Primary OPC culture

Mice brains from P4-P6 pups were removed, dissociated (see above) and the single cell suspension was used to enrich for OPCs by CD140a microbeads (Miltenyi Biotec, 130-101-502) according to manufacturer’s instructions. Brains from 4-5 mice were pooled for one batch of OPC culture. Cells were seeded on petri dishes or multi-well plates pre-coated with poly-L lysine (Sigma, P4707) for >1h at 37C, and Fibronectin for >1h (Sigma, F1141, 1mg/ml stock, 1:1000 diluted in 1x PBS). Cells were cultivated in OPC proliferation medium -DMEM/Gmax (ThermoFisher Scientific, 10565018), 1x N2 supplement (ThermoFisher Scientific, 17502048), 1x penicillin–streptomycin (ThermoFisher Scientific, 15140122), 1xNeuroBrew (Miltenyi 130-093-566), bFGF 20 ng ml−1 (Peprotech, 100-18B) and PDGF-AA 10 ng ml−1 (Peprotech, 100-13A) until confluency (5-6 days), passaged once and collected 72h later.

### HiChIP

HiChIP was performed in three biological replicates with at least 1 milion cells used as input. Briefly, cultured OPCs were collected using TrypLE Express (Gibco, 12604013), washed once with 10ml of 1xPBS and crosslinked using freshly prepared 1% formaldehyde (Methanol-free, Pierce, 28906) diluted in 1x PBS for 15 minutes at room temperature with gentle rotation. Formaldehyde was quenched by addition of final 125 mM glycine and incubated for 5 min at room temperature with gentle rotation. Fixed cells were then washed once with ice cold 1x PBS, flash frozen and stored at -80 until processed. HiChIP was performed as in Mumbach et al^31^. Briefly, chromatin was sonicated using the covaries ME220 with settings 75 PIP, 5% duty cycle, 200 cycles/burst for 2 minutes (for 1-3 milion cells). The immunoprecipitation was performed using 2 ug of H3K27ac antibody (Abcam, Ab177178) and 20 ul of protein A dynabeads (Thermo Fisher, 007613560), with 0.75 ul of in-house produced Tn5 used for tagmentation and 15-16 cycles of final PCR amplification (NEBNext High Fidelity 2x PCR mastermix, M0541L). Barcoded libraries wer gel-purified, quantified using bioanalyzer, mixed in equimolar ratios and sequenced on one flow cell of NovaSeq S Prime (Illumina).

### HiChIP data processing

Paired-end sequencing reads from HiChIP experiments were aligned to mm10 genome and processed using the HiC-Pro pipeline^40^. The pipeline’s hicpro2juicebox.sh script was used to generate .hic files which were loaded into Juicebox^41^ for viewing contact maps. The hic2cool tool (https://github.com/4dn-dcic/hic2cool) was used to generate 5kb-resolution .cool files.

### ABC model

ATAC-seq data was downloaded from 10x genomics online resources (https://support.10xgenomics.com/single-cell-atac/datasets/1.2.0/atac_v1_adult_brain_fresh_5k) and cell type specific peaks were called using MACS2. Cell-type specific H3K27ac bam files were generated from the cellranger-ATAC output files. Gene expression data from the scRNA-seq was used to identify the candidate genes and high-resolution Hi-C data from neural progenitor cells^30^ was used to estimate contact frequency. Default parameters were used for generating the candidate enhancer list. Regulatory loops with an ABC score > 0.02 were used for downstream analyses.

### Pileup analysis

The coolpuppy package was used for generating pileup profiles^42^. In brief, OLG-specific enhancer-promoter BEDPE files from the ABC model were used as pre-defined candidate regions to look for enrichment. Pileups were performed on OPC-derived H3K27ac HiChIP data and mESC data^31^. All HiChIP data was balanced using the cooler library prior to performing the pileups^43^.

## Acknowledgements

We would like to thank Tony Jimenez-Beristain for writing laboratory animal ethics permit 1995_2019 and assistance with animal experiments, Leslie Kirby, Mandy Meijer and Petra Kukanja for assistance with animal experiments, Bastien Hervé for setting up the Shiny web resource, Simon Elsasser and Rickard Sandberg for proofreading and providing critical comments on the manuscript, Ida Johansson and Elin Dunevall from Karolinska Institute Protein Science facility for cloning the pA-Tn5 construct and the protein purification, Matilda Eriksson for performing the 10x ATAC-seq protocol, the Single Cell Genomics Facility and the staff at Comparative Medicine-Biomedicum. The authors acknowledge support from the National Genomics Infrastructure in Stockholm funded by Science for Life Laboratory, the Knut and Alice Wallenberg Foundation and the Swedish Research Council, and Swedish National Infrastructure for Computing /Uppsala Multidisciplinary Center for Advanced Computational Science for assistance with massively parallel sequencing and access to the UPPMAX computational infrastructure. M.B. is funded by the Vinnova Seal of Excellence Marie-Sklodowska Curie Actions grant RNA-centric view on Oligodendrocyte lineage development (RODent). Work in G.C.-B.’s research group was supported by the Swedish Research Council (grant 2015-03558 and 2019-01360), the European Union (FP7/Marie Curie Integration Grant EPIOPC, Horizon 2020 Research and Innovation Programme/European Research Council Consolidator Grant EPIScOPE, grant agreement number 681893), the Swedish Brain Foundation (FO2017-0075), the Ming Wai Lau Centre for Reparative Medicine, the Swedish Cancer Society (Cancerfonden; CAN2016/555), Knut and Alice Wallenberg Foundation (grant 2019-0107), The Swedish Society for Medical Research (SSMF, grant JUB2019), Ming Wai Lau Centre for Reparative Medicine and Karolinska Institutet.

## Data availability

Raw data is deposited in GEO XXXXX and code is available at https://github.com/Castelo-Branco-lab/scCut-Tag_2020. The scCut&Tag dataset can be explored at the following web resource https://castelobranco.shinyapps.io/BrainCutAndTag2020/

## Author Contributions

MB and GCB conceived the study, designed the experiments and analysis and wrote the manuscript. MB optimized and performed the scC&T experiments and analyzed the scC&T data. MK and MB performed the HiChIP experiment. MK analyzed the HiChIP data and helped with generation of related figures. All authors contributed and approved the manuscript.

## Supplementary Tables

Supplementary table 1. List of marker regions (bins) defined by the scC&T data.

Supplementary table 2. List of enhancer promoter loops predicted by ABC model.

Supplementary table 3. List of correlated H3K27ac peak regions predicted by cicero.

## Figure legends

**Extended data figure 1.**
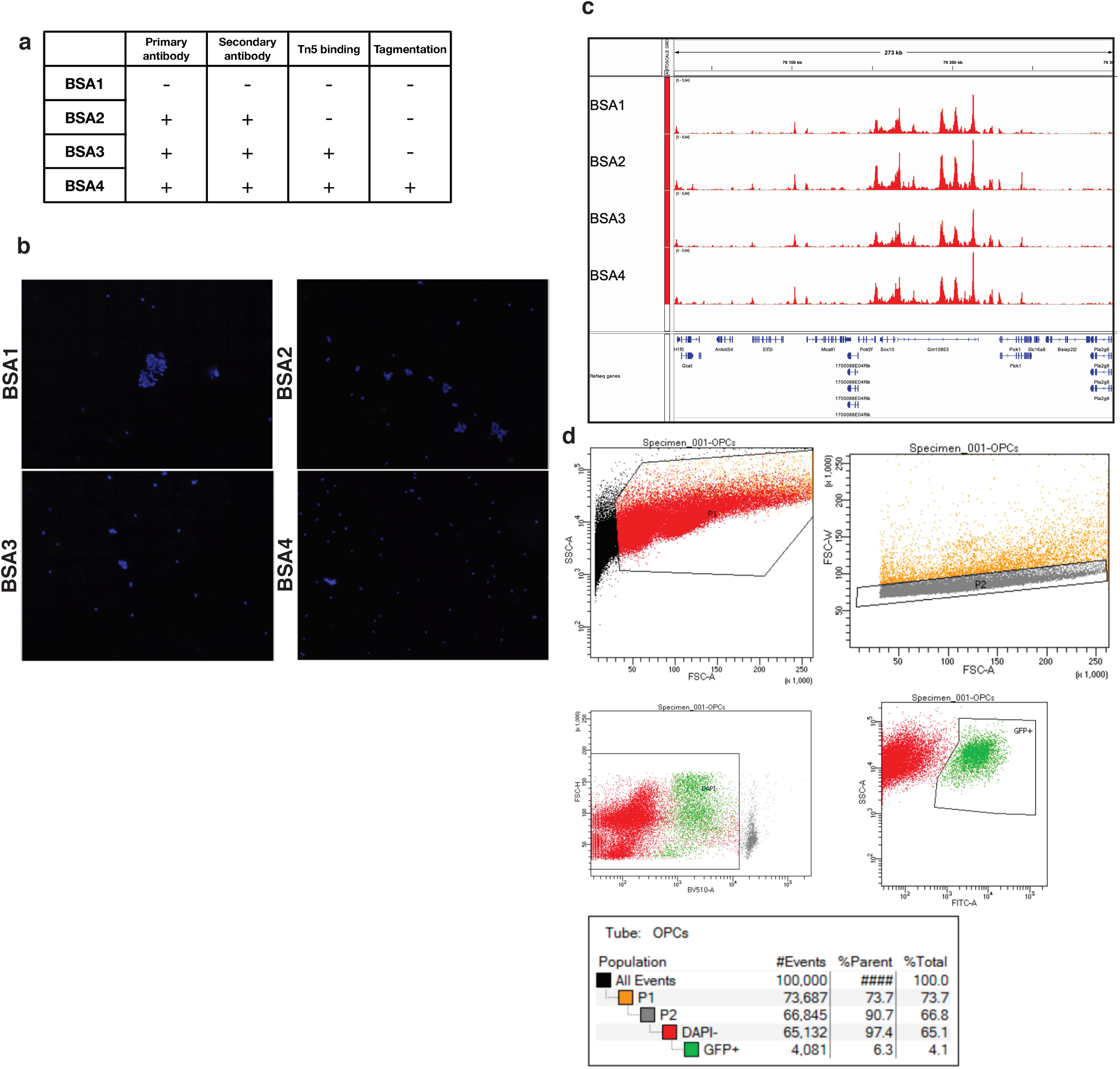
Optimization of tagmentation step in the scC&T protocol. a. Table depicting scC&T protocol steps (Primary Antibody, Secondary Antibody, Tn5 binding, Tagmentation) and whether BSA was included in the scC&T buffers during these steps. b. DAPI counterstaining of the nuclei after the scC&T procedure. Inclusion of BSA in the procedure significantly reduces clumping of the nuclei. c. Genome browser profiles of bulk cut&Tag experiment from the different BSA conditions (described in a). d. Gating strategy depicting sorting of GFP cells. P1 Depicts general gate for selection of cells, P2 gate selects for singlets, DAPI-gate selects only live cells and GFP+ gate selects cells that posess GFP signal (Sox10-Cre/GFP).

**Extended data figure 2.**
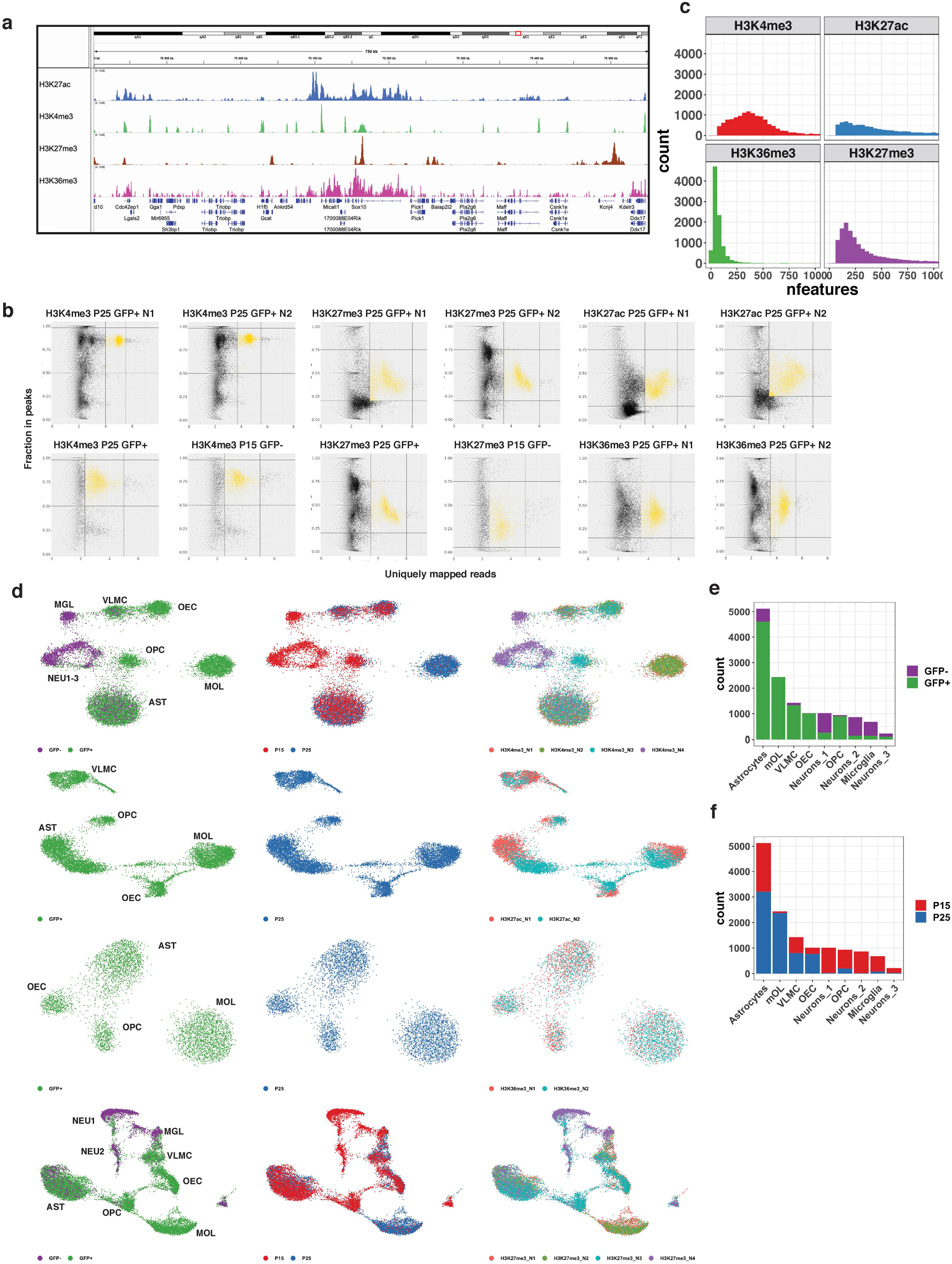
Quality control of the scC&T data. a. Merged pseudobulk profiles of scC&T with four antibodies against modified histones. b. Scatterplot of number of reads per cells (x axis) and fraction of reads originating from peak regions (y axis) that was used to set cutoffs for cells identification. Cut-offs were set after manual inspection of the plots and are depicted as horizontal and vertical lines overlaid over the plot. c. Histogram of number of features identified per cell for each antibody used in scC&T. d. Two dimensional UMAP embedding of the scC&T data. Cells are colored by correspondence to GFP population, developmental age and biological replicate. e-f. Barplot summary of the correspondence to the (e) GFP population and (f) developmental age per cell type identified from the H3K4me3 sec&T data.

**Extended data figure 3.**
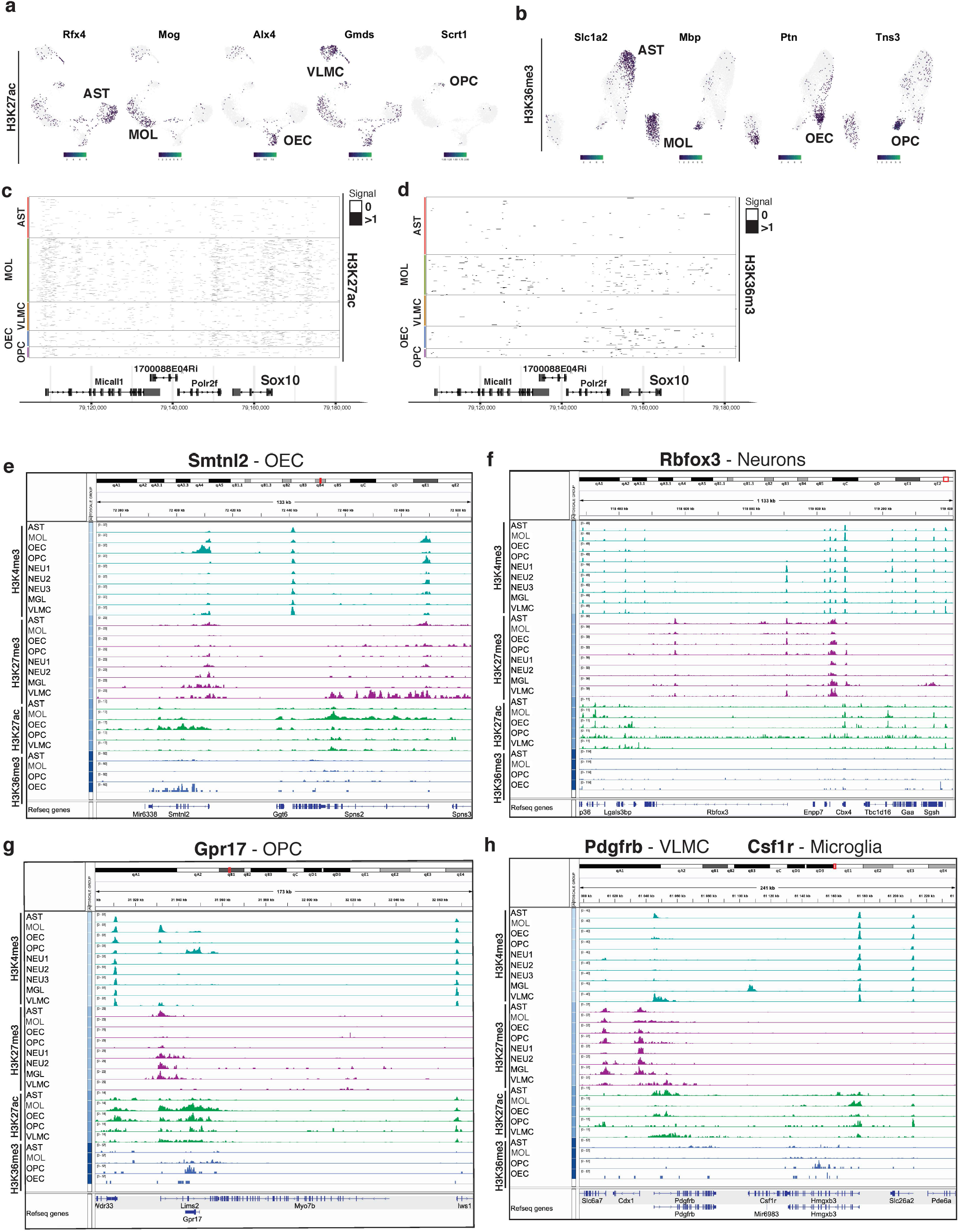
De nova identification of cell types by cell type specific marker regions. Projection of gene activity scores of (a) H3K27ac and (b) H3K36me3 scC&T on the two-dimensional UMAP embedding. Gene name is depicted in the title and specific population is highlighted in the UMAP plot by labeling the cell type. c-d. Heatmap representation of the scC&T signal for (c) H3K27ac and (d) H3K36me3. X axis represents genomic region, each row in Y axis contains data from one cell. Cell correspondence to clusters is depicted by color bar on the right side of the heatmap and annotated with cell type. Signal is aggregated per 250 bp windows and binarized. e-h. Aggregated pseudobulk scC&T profiles for four histone modifications in all identified cell types around selected marker genes

**Extended data figure 4.**
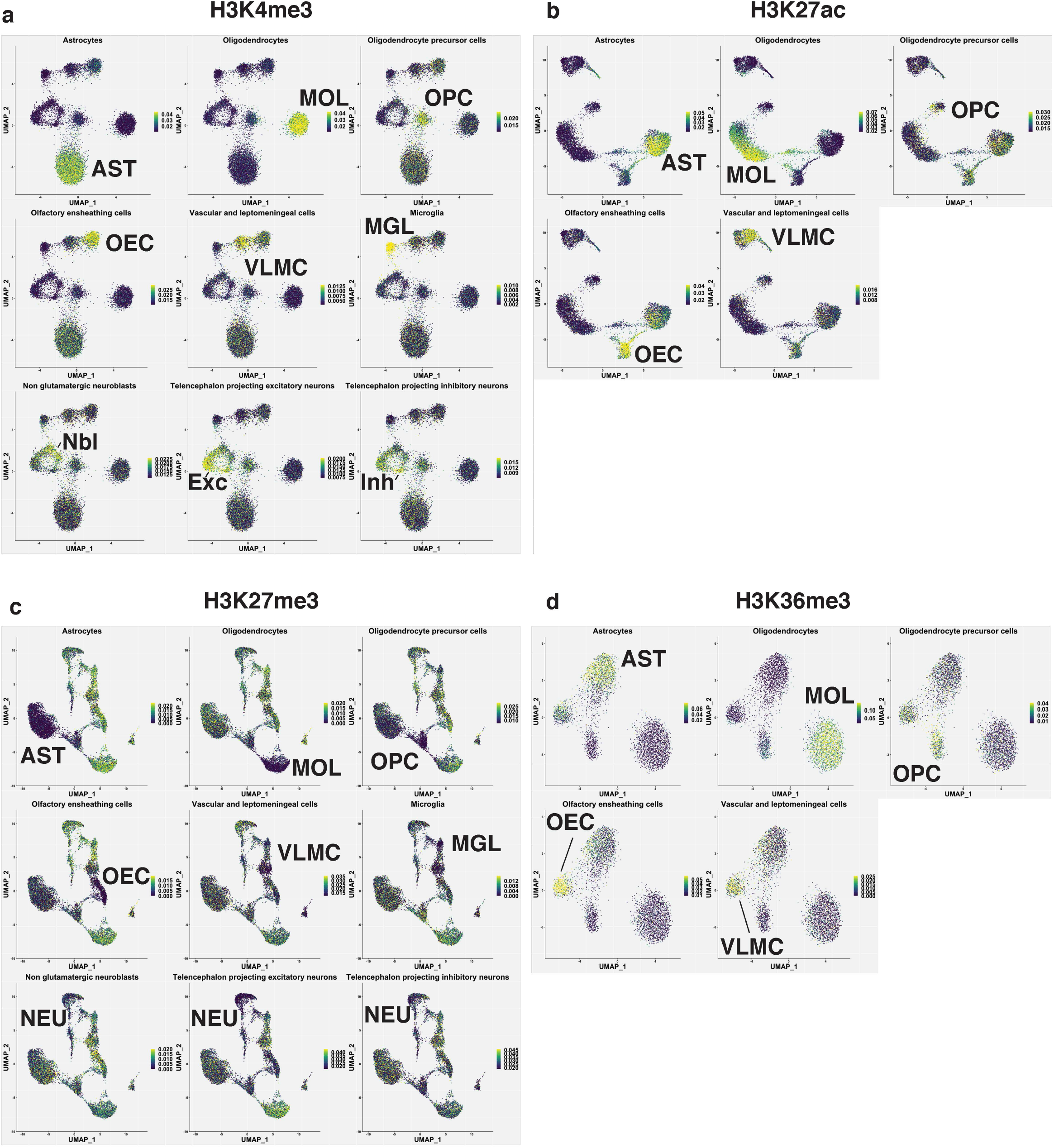
Metagene analysis of gene activity scores. a-d. Metagene activity projection of scC&T data on the UMAP embeddings of four histone modification scC&T datasets. Metagenes are selected as top 100 most specifically expressed in the scRNA-seq data3.

**Extended data figure 5.**
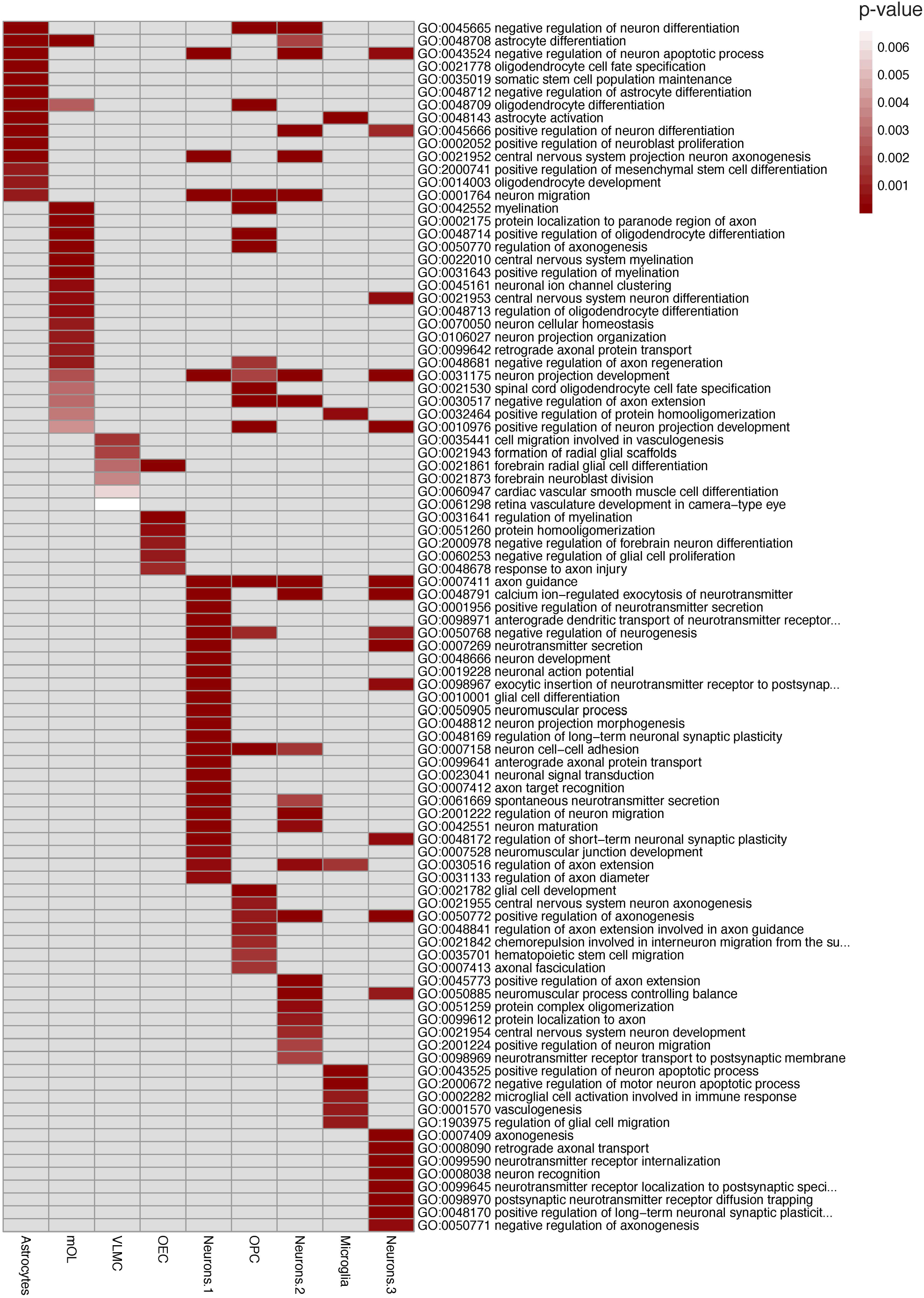
GO analysis. a. Gene ontology analysis of the marker genes determined by gene activity scores from the H3K4me3 scC&T data. GO terms were manually selected from the list of all enriched GO terms in all populations.

**Extended data figure 6.**
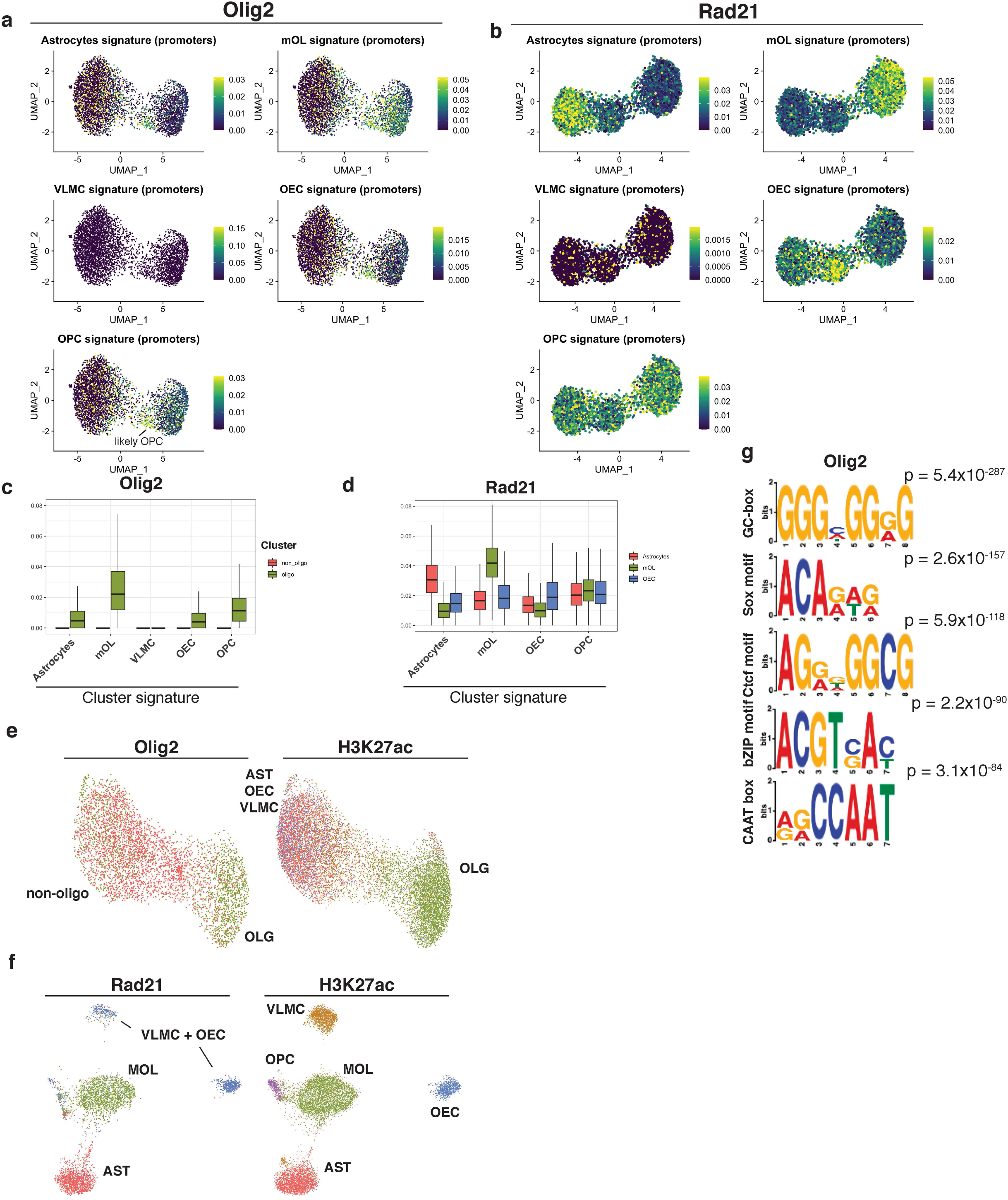
scC&T of transcription factors. a. Meta-region activity scores of marker regions determined from H3K4me3 scC&T data and specific for the respective population for (a) Olig2 and (b) Rad21 scC&T data. c-d. Boxplot representation of a and b, single cell meta-region profiles aggregated per cell type. e-f. Co-embedding of (e) H3K27ac and Olig2 and (f) H3K27ac and Rad21 in single two dimensional UMAP space. g. Additional motifs identified using MEME from the merged pseudobulk profile of Olig2 scC&T.

## References

1. Skene, P. J. & Henikoff, S. An efficient targeted nuclease strategy for high-resolution mapping of DNA binding sites. eLife 6,.

2. Kaya-Okur, H. S. et al. CUT&Tag for efficient epigenomic profiling of small samples and single cells. Nat. Commun. 10, 1930 (2019).

3. Zeisel, A. et al. Molecular Architecture of the Mouse Nervous System. Cell 174, 999-1014.e22 (2018).

4. La Manno, G. et al. RNA velocity of single cells. Nature 560, 494–498 (2018).

5. Buenrostro, J. D. et al. Single-cell chromatin accessibility reveals principles of regulatory variation. Nature 523, 486–490 (2015).

6. Lorthongpanich, C. et al. Single-Cell DNA-Methylation Analysis Reveals Epigenetic Chimerism in Preimplantation Embryos. Science 341, 1110–1112 (2013).

7. Rotem, A. et al. Single-cell ChIP-seq reveals cell subpopulations defined by chromatin state. Nat. Biotechnol. 33, 1165–1172 (2015).

8. Ai, S. et al. Profiling chromatin states using single-cell itChIP-seq. Nat. Cell Biol. 21, 1164–1172 (2019).

9. Wang, Q. et al. CoBATCH for High-Throughput Single-Cell Epigenomic Profiling. Mol. Cell 76, 206-216.e7 (2019).

10. Ku, W. L. et al. Single-cell chromatin immunocleavage sequencing (scChIC-seq) to profile histone modification. Nat. Methods 16, 323–325 (2019).

11. Carter, B. et al. Mapping histone modifications in low cell number and single cells using antibody-guided chromatin tagmentation (ACT-seq). Nat. Commun. 10, 3747 (2019).

12. Falcão, A. M. et al. Disease-specific oligodendrocyte lineage cells arise in multiple sclerosis. Nat. Med. 24, 1837–1844 (2018).

13. Marques, S. et al. Transcriptional Convergence of Oligodendrocyte Lineage Progenitors during Development. Dev. Cell 46, 504-517.e7 (2018).

14. Jäkel, S. et al. Altered human oligodendrocyte heterogeneity in multiple sclerosis. Nature 566, 543–547 (2019).

15. Marques, S. et al. Oligodendrocyte heterogeneity in the mouse juvenile and adult central nervous system. Science 352, 1326–1329 (2016).

16. Sousa, V. H., Miyoshi, G., Hjerling-Leffler, J., Karayannis, T. & Fishell, G. Characterization of Nkx6-2-Derived Neocortical Interneuron Lineages. Cereb. Cortex 19, i1–i10 (2009).

17. Matsuoka, T. et al. Neural crest origins of the neck and shoulder. Nature 436, 347–355 (2005).

18. Huang, W. et al. Novel NG2-CreERT2 knock-in mice demonstrate heterogeneous differentiation potential of NG2 glia during development. Glia 62, 896–913 (2014).

19. Zhu, X. et al. Age-dependent fate and lineage restriction of single NG2 cells. Development 138, 745–753 (2011).

20. Zhu, X., Bergles, D. E. & Nishiyama, A. NG2 cells generate both oligodendrocytes and gray matter astrocytes. Development 135, 145–157 (2008).

21. Stuart, T. et al. Comprehensive Integration of Single-Cell Data. Cell 177, 1888-1902.e21 (2019).

22. Wang, J. et al. EED-mediated histone methylation is critical for CNS myelination and remyelination by inhibiting WNT, BMP, and senescence pathways. Sci. Adv. 6, eaaz6477 (2020).

23. Benayoun, B. A. et al. H3K4me3 Breadth Is Linked to Cell Identity and Transcriptional Consistency. Cell 158, 673–688 (2014).

24. MEME Suite: tools for motif discovery and searching. https://www.ncbi.nlm.nih.gov/pmc/articles/PMC2703892/.

25. Li, Y. et al. The structural basis for cohesin–CTCF-anchored loops. Nature 578, 472–476 (2020).

26. Yu, Y. et al. Olig2 Targets Chromatin Remodelers To Enhancers To Initiate Oligodendrocyte Differentiation. Cell 152, 248–261 (2013).

27. Darr, A. J. et al. Identification of genome-wide targets of Olig2 in the adult mouse spinal cord using ChIP-Seq. PLoS ONE 12, (2017).

28. Wißmüller, S., Kosian, T., Wolf, M., Finzsch, M. & Wegner, M. The high-mobility-group domain of Sox proteins interacts with DNA-binding domains of many transcription factors. Nucleic Acids Res. 34, 1735–1744 (2006).

29. Fulco, C. P. et al. Activity-by-contact model of enhancer–promoter regulation from thousands of CRISPR perturbations. Nat. Genet. 51, 1664–1669 (2019).

30. Bonev, B. et al. Multiscale 3D Genome Rewiring during Mouse Neural Development. Cell 171, 557-572.e24 (2017).

31. Mumbach, M. R. et al. Enhancer connectome in primary human cells identifies target genes of disease-associated DNA elements. Nat. Genet. 49, 1602–1612 (2017).

32. Pliner, H. A. et al. Cicero Predicts cis-Regulatory DNA Interactions from Single-Cell Chromatin Accessibility Data. Mol. Cell 71, 858-871.e8 (2018).

33. Moore, J. E. et al. Expanded encyclopaedias of DNA elements in the human and mouse genomes. Nature 583, 699–710 (2020).

34. Zhang, Y. et al. Model-based analysis of ChIP-Seq (MACS). Genome Biol. 9, R137 (2008).

35. Street, K. et al. Slingshot: cell lineage and pseudotime inference for single-cell transcriptomics. BMC Genomics 19, 477 (2018).

36. Khan, A. et al. JASPAR 2018: update of the open-access database of transcription factor binding profiles and its web framework. Nucleic Acids Res. 46, D260–D266 (2018).

37. Ramírez, F. et al. deepTools2: a next generation web server for deep-sequencing data analysis. Nucleic Acids Res. 44, W160–165 (2016).

38. Wickham, H. ggplot2: Elegant Graphics for Data Analysis. (Springer-Verlag, 2009). doi: 10.1007/978-0-387-98141-3.

39. Köster, J. & Rahmann, S. Snakemake—a scalable bioinformatics workflow engine. Bioinformatics 28, 2520–2522 (2012).

40. Servant, N. et al. HiC-Pro: an optimized and flexible pipeline for Hi-C data processing. Genome Biol. 16, 259 (2015).

41. Durand, N. C. et al. Juicebox Provides a Visualization System for Hi-C Contact Maps with Unlimited Zoom. Cell Syst. 3, 99–101 (2016).

42. Flyamer, I. M., Illingworth, R. S. & Bickmore, W. A. Coolpup.py: versatile pile-up analysis of Hi-C data. Bioinformatics 36, 2980–2985 (2020).

43. Abdennur, N. & Mirny, L. A. Cooler: scalable storage for Hi-C data and other genomically labeled arrays. Bioinformatics 36, 311–316 (2020).

